# *Hallucigenia*’s diet illuminates the feeding ecology of Cambrian lobopodians

**DOI:** 10.64898/2025.12.28.696761

**Authors:** Javier Ortega-Hernández

## Abstract

The armoured lobopodian *Hallucigenia sparsa* embodies the seemingly uncanny nature of the animals that evolved during the Cambrian Explosion over 500 million years ago. Initially regarded as an evolutionary oddball, the exceptional preserved anatomy of *H. sparsa* has been substantially revised, leading to a better understanding of its relationships with other lobopodians and phylogenetic affinities with extant panarthropod phyla. However, the ecology and behaviour of *H. sparsa* remain largely enigmatic owing to the difficulties of interpreting its functional morphology and the perceived lack of modern analogues. Restudy of a composite fossil assemblage from the middle Cambrian Burgess Shale demonstrates swarm-like behaviour of several small *H. sparsa* individuals scavenging on a dead ctenophore. The lack of grasping, masticatory, or piercing mouthparts in *H. sparsa* points to suction feeding as a viable strategy to consume the gelatinous carcass. Reassessment of the morphology of *H. sparsa* reveals functional analogues shared with extant pycnogonids, including an elongate anterior end with a terminal mouth opening and enlarged foregut chambers lined up with sclerotized denticles. These observations suggest that suction feeding is ancestral for stem-group Onychophora and highlights the critical trophic role of small-bodied armoured lobopodians as degraders of soft-bodied carcasses in Cambrian benthic ecosystems.

## Introduction

Cambrian sites of exceptional preservation have produced critical insights into the autecology, interactions, and life strategies of early animals [1]. The feeding behaviour of Cambrian animals is typically inferred through functional interpretations of the fossilized morphology, such as complex mouthparts [02, 03, 04], or specialized post-oral appendages [05, 06, 07] and internal organ systems [08, 09]. More rarely, it is possible to infer feeding preferences based on gut contents or preserved interspecific fossil assemblages [1, 10, 11]. However, several taxa lack discrete adaptations that would inform their feeding ecology, either due to preservation biases, because the morphology is uninformative and/or difficult to interpret, or because they lack clear modern analogues. These challenges not only affect our understanding of individual species’ ecology, but also fundamentally hinder efforts to accurately reconstruct marine trophic networks during the Cambrian and thus misinform our view of the ecological complexity of the earliest bilaterian dominated marine communities [12, 13]. Among Cambrian lobopodians – paraphyletic leg-bearing worm-like ecdysozoans that include early branching members of all extant panarthropod phyla [14] – the feeding ecology of hallucigeniids and other small bodied armoured forms is poorly known [15, 16, 17]. Whereas some Cambrian lobopodians feature elaborate limbs for suspension feeding [05, 06] or complex digestive glands for processing food items [08], hallucigeniids have comparatively modest morphologies with a simple bulbous head and up to three sets of claw-less anterior tentacles [16, 18, 19, 20]. Sclerotized mouthparts for hallucigeniids have only been described in the mid-Cambrian Burgess Shale *Hallucigenia sparsa*, which consist of a buccal cavity, a ring of internal circumoral elements and a row of acicular foregut denticles, whereas the digestive system is a simple gut tract [18]. Despite this data and the fact that *H. sparsa* represents the most emblematic lobopodian and one of the best known soft-bodied Cambrian organisms, details of its feeding ecology and behaviour remain largely unknown. Restudy of a unique composite specimen reveals the only known evidence of collective scavenging behaviour in *H. sparsa* offering new insights on the feeding behaviour and lifestyle of this iconic species. The findings have direct implications for better understanding the enigmatic ecology of armoured lobopodians 15, 16, 17, and provide insights into the processes that facilitated nutrient cycling of Cambrian benthic ecosystems.

## Results

The studied material (Figs. 1, 2; Figs. S1, S2A), MCZ.IP.102104 (previously MCZ 1084), is a complex assemblage that was first illustrated in the original systematic description of *H. sparsa* in an upside-down orientation [21]. The specimen has been subsequently referred to as the only direct evidence of scavenging in *H. sparsa* [15, 17], but not formally restudied in over 45 years. The fossil originates from the mid-Cambrian (Wuliuan) Burgess Shale in British Columbia, more specifically from a stratigraphic level approximately 70 feet above Walcott Quarry in Mount Stephen. First regarded as the remains of a single new unidentified worm carcass associated with several *H. sparsa* individuals [21], re-examination of MCZ.IP.102104 reveals unnoticed details of its affinities and broader ecological significance. The carcass (width ca. 35 mm, height ca. 19 mm) corresponds to a soft-bodied organism with a labile cellular body wall based on the preservation of two-dimensional carbonaceous films, including both dark areas and highly reflective patches. There is evidence of substantial distortion caused by decay, namely the presence of irregular margins and rupture between the larger highly reflective area on the bottom (Fig. 1A, 2A) relative to a smaller but morphologically complex darker top area of the carcass. The specimen is composed of a series of serially repeating carbonaceous parallel strands spaced at irregular intervals, including at least a dozen strands on the reflective bottom area, at least six thick strands on the darker top area, and with one strand directly connecting the two regions (arrowhead in Fig. 1B, C, 2B). Critically, the six thick strands on the darker top area and the single strand connected to the reflective bottom area show a radial organization and converge into a well-defined ring-shaped structure surrounded by thinner strands and located approximately at the mid-point of the specimen’s width (Fig. 1B, C, H). Comparative anatomy indicates that the carcass belongs to a pelagic ctenophore (comb jelly) [22, 23] (Fig. S2). The evenly spaced parallel strands with a radial organization represent the comb rows, the ring-shaped structure is the apical organ that defines the aboral region, and the thin strands associated with the former correspond to ciliary grooves. The carcass represents a complete individual preserved in slightly oblique aboral view that has become almost entirely bisected on the transverse plane, with the two parts dangling together from one comb row. The presence at least 18 observable comb rows suggests that their total number was higher, and thus comparable with the morphology of the Burgess Shale stem-group ctenophores *Ctenorhabdotus* and *Xanioascus*, both of which bear 24 comb rows and are also known from Mount Stephen sites in the Burgess Shale [22]. The seeming lack of a capsule on the apical organ (diagnostic of *Ctenorhabdotus* [23]), slender comb rows, overall sac-like appearance of the carcass, and possible presence of internal ovoid bodies suggest a tentative assignment to *Xanioascus* [22] (Fig. S2).

**Figure 1.**
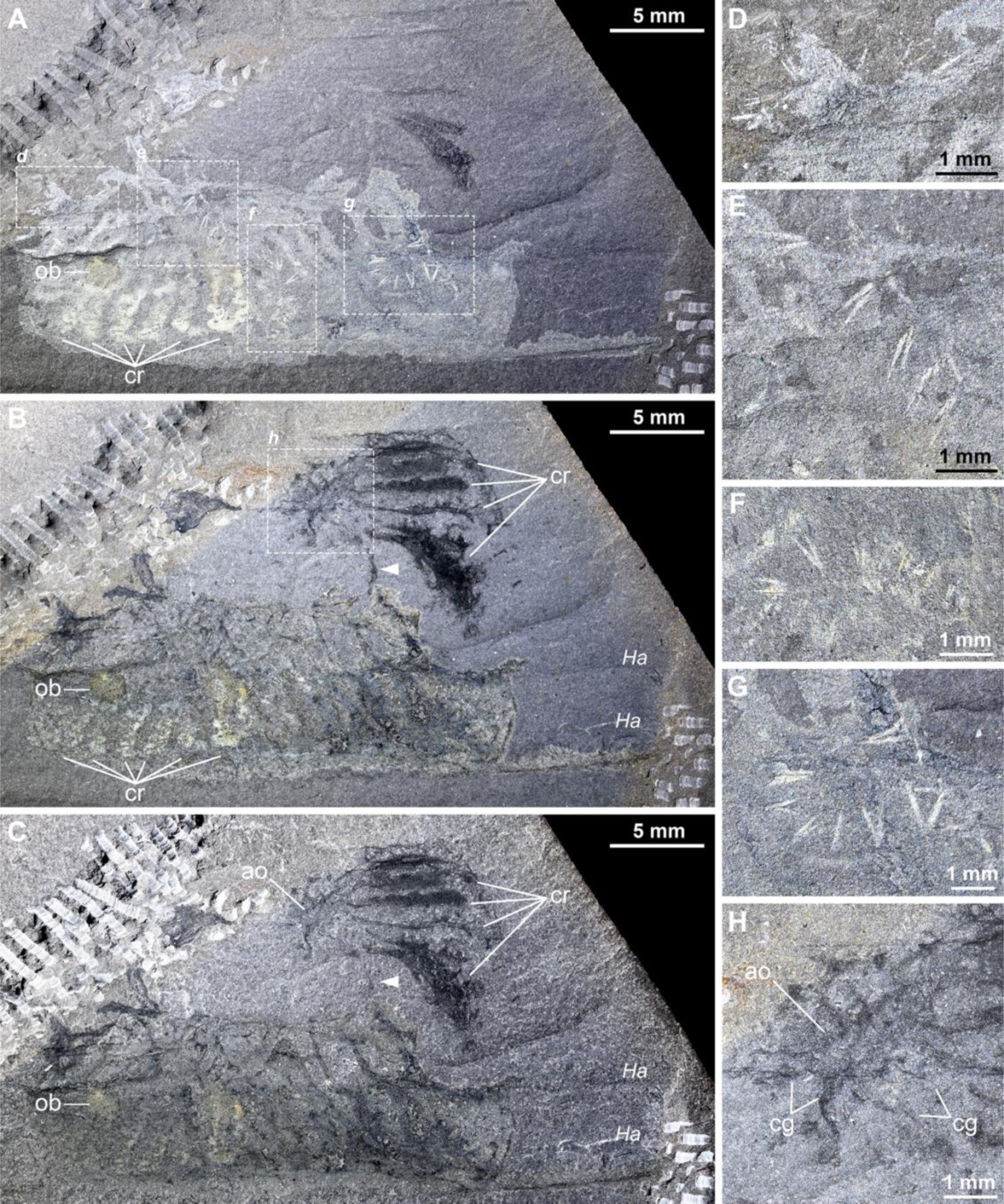
Fossil assemblage part showing seven small individuals of *Hallucigenia sparsa* feeding on carcass of the ctenophore *Xanioascus* from the mid-Cambrian Burgess Shale. **A.** MCZ.IP.102104 part, photographed wet under cross polarized illumination emphasizing highly reflective bottom area with comb rows and five *H. sparsa* individuals. **B.** MCZ.IP.102104 part, photographed wet under cross polarized illumination emphasizing dark top area with apical organ and ciliary grooves and single comb row connecting both parts of the specimen (arrowhead). **C.** MCZ.IP.102104 part, photographed dry under cross polarized illumination. **D-G.** Magnification of subpanels in A showing complete small individuals of *H. sparsa*. **H.** Magnification of subpanel in B showing details of the apical organ and ciliary grooves. Abbreviations. ao, apical organ; cg, ciliary grooves; cr, comb rows; *Ha*, isolated *Hallucigenia* spines; ob, ovoid body.

**Figure 2.**
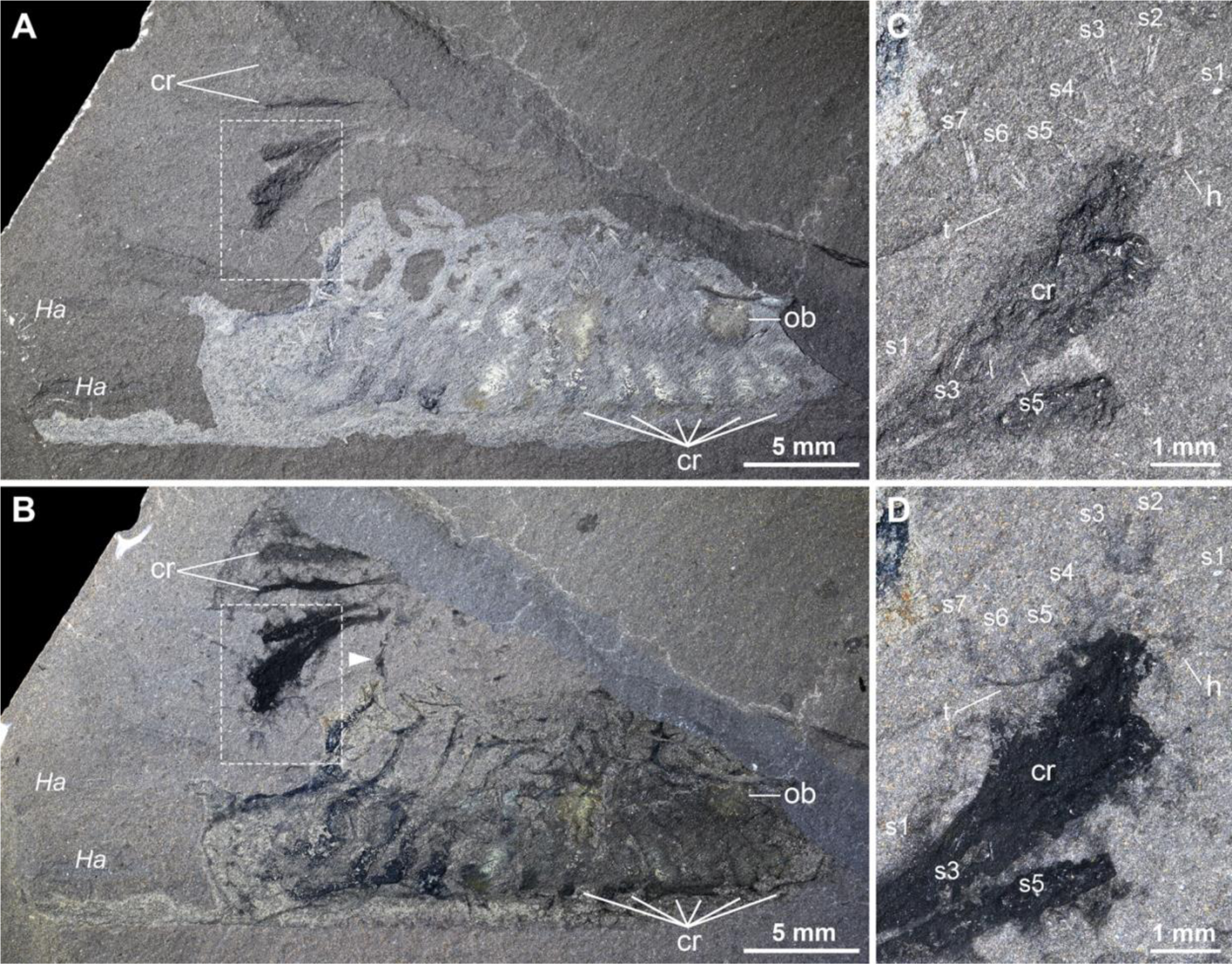
Fossil assemblage counterpart showing seven small individuals of *Hallucigenia sparsa* feeding on carcass of the ctenophore *Xanioascus* from the mid-Cambrian Burgess Shale. **A.** MCZ.IP.102104 counterpart, photographed wet under cross polarized illumination emphasizing highly reflective areas. **B.** MCZ.IP.102104 counterpart, photographed wet under cross polarized illumination emphasizing dark areas including single comb row connecting both parts of *Xanioascus* carcass (arrowhead). **C.** Magnification of subpanel in A (rotated clockwise 180 degrees) showing well preserved *H. sparsa* individuals with complete set of spines and soft tissues. **D.** Magnification of subpanel in B (rotated clockwise 180 degrees). Abbreviations: cr, comb rows; h, head; Ha, isolated *Hallucigenia* spines; ob, ovoid body; s*n,* dorsal spines; t, trunk.

The uniqueness of MCZ.IP.102104 consists of the presence of several diminutive individuals of *H. sparsa* directly associated with the *Xanioascus* carcass. An estimate of 18 specimens were initially reported [21], but close examination confirms the presence of seven well preserved complete *H. sparsa* individuals (Fig. 1D-H, 2C, D), with additional isolated spine pairs scattered across the slab. All complete *H. sparsa* individuals have a body length size range of ca. 3-6mm, making them the smallest representatives of this species known from articulated remains [18], and also the smallest hallucigeniids reported to date [19, 20, 24] (Fig. S3). Five evenly spread-out *H. sparsa* can be observed on the reflective bottom area of the *Xanioascus* carcass (Fig. 1A, D-G), and the two smallest ones associated with the dark top area that contains the apical organ and ciliary grooves. The latter bears a particularly well-preserved individual with a total length of ca. 3.5 mm featuring a complete dorsal set of seven pairs of spines as well as delicate traces of the head and trunk regions (Fig. 2C, D).

## Discussion

### Taphonomic interpretation

MCZ.IP.102104 reveals new insights into the feeding ecology and lifestyle of *Hallucigenia sparsa* thanks to an improved understanding of the overall morphology and evolutionary significance of this armoured lobopodian since its initial publication [18, 25], as well as a more comprehensive view of animal biodiversity preserved in Cambrian Burgess Shale-type deposits [22, 23]. The *Xanioascus* carcass shows clear signs of post-mortem alteration including the rupture of the body wall separating it into two fragments and distortion of the original morphology due to compression and the oblique angle of burial. Considering the extremely labile constitution of the gelatinous body wall of pelagic ctenophores, the *Xanioascus* carcass most likely rapidly settled on the seafloor after death. The fact that the delicate comb rows and ciliary grooves maintain a substantial degree of overall integrity and their relative organization despite the rupturing of the body wall suggest that the carcass likely stabilized in a low energy local depositional environment with restricted oxygen. These conditions would have reduced the rate of fragmentation and aerobic decay [26] and deterred medium and large (ca. body length > 1 cm) carnivorous scavengers reliant on well oxygenated conditions [27]. The preservation quality of the *Xanioascus* carcass contrasts with that of the seven *H. sparsa* individuals, all of which are completely articulated based on the organization of the dorsal spines despite their small size (Fig. 1D-G), and even show remains of the soft-bodied trunk and head (Fig. 2C, D). *H. sparsa* specimens show no indication of decay, breakage, or alteration, indicating that they were most likely still alive at the time of burial. The differences in preservation quality between *Xanioascus* and *H. sparsa,* and the fact that all complete *H. sparsa* individuals are in direct contact with the ctenophore carcass support the interpretation that this fossil assemblage reflects a legitimate ecological interaction, and suggest rapid burial *in situ* with minimal or no transport.

### Ecology and feeding behaviour of Hallucigenia sparsa

The capricious morphology of hallucigeniids has prompted speculation about their ecology and overall mode of life. Although there is a broad intuitive agreement that the armature of dorsal spines in hallucigeniids and other armored lobopodians had a defensive function against potential predators [05, 06, 14, 16], and likely also served as sites of muscle attachment and limb locomotion leverage [28], other aspects of their lifestyle are less resolved. Cambrian trophic network studies have repeatedly referenced the supposed co-occurrence between *Hallucigenia sparsa* and the sponge *Vauxia* [12, 13] and assumed a feeding interaction. This idea appears to originate from a brief statement in the original description speculating “it is possible that *H. sparsa* included sessile creatures such as sponges in its diet” [21]. However, the supposed association between these taxa has never been investigated in detail and remains anecdotal at best, so there is no direct evidence that *H. sparsa* specifically interacted with *Vauxia*, much less fed on it or any other sponge for that matter. Interpreting the functional morphology of the oral structures of *H. sparsa* is not straightforward due to their uniqueness [18], and further challenged by the simple digestive tract and complete absence of gut contents in this species, and hallucigeniids more broadly. It has also been hypothesized that hallucigeniids could be epibenthic suspension feeders [29]; however, this interpretation rests on phylogenetic inference alone, and is not directly supported by the functional morphology of hallucigeniid lobopodians as they lack setulose limbs or other similar adaptations typically associated with this feeding strategy.

MCZ.IP.102104 represents the best evidence of feeding behaviour in *H. sparsa* to date, and the association with *Xanioascus* offers valuable insights on its ecology beyond its interpretation as a simple scavenger [21, 15, 17]. First, it shows that *H. sparsa* fed on gelatinous carcasses, which would be most likely achieved by suctioning body fluids using its elongate bulbous head with a terminal mouth. This observation can be reasonably generalized to an ability to scavenge on soft-bodied carcasses with cellular body walls, such as medusozoans [30], shell-less mollusks [31] or early vertebrates [32], but it is impossible to assess the degree of selectivity of food preference with the available data. An interpretation for suction feeding in *H. sparsa* is consistent with the presence of the enlarged buccal chamber [18] (Fig. 3B, C) and the lack of external adaptations for food processing such as raptorial appendages, teeth or scalids needed for a macrophagous diet as observed in large-bodied lobopodians [08, 15]. In this context, the presence of sclerotized structures in the foregut, including the ring of circum-oral elements and the row aciculae-like denticles, may have functioned as a rudimentary internal filtering system capturing ingested solid particles based on comparisons with extant marine euarthropods with a suction feeding strategy (see below) [33]. A fluid-based diet would also explain the absence of complex digestive glands in the gut tract of *H. sparsa*, otherwise commonplace in macrophagous Cambrian lobopodians [08], and non-existent gut contents despite the availability of dozens of carefully studied specimens [18]. The three pairs of claw-less tentacle-like anterior lobopodous appendages of *H. sparsa* are well suited for a sensorial function to locate appropriate food items, whereas the claw-bearing walking legs are well adapted for anchoring and locomotion (Fig. 3A). Thus, although it has been considered that there are no obvious modern counterparts for hallucigeniids based on their peculiar overall morphology [15], these observations suggest that pycnogonids (sea spiders) might be a sensible ecological analogue. Pycnogonids are slow moving epibenthic marine chelicerates that have tubular bodies and typically elongate clawed appendages used for walking and anchoring to various substrates (Fig. 3E). Pycnogonid mouthparts boast disparate morphological adaptations to consume diverse food items (e.g. algae, sponges, cnidarians, bilaterians), with the main common denominator being that they perform suction feeding of the body fluids using their muscular anterior proboscis [33]. Although pycnogonids typically have paired raptorial appendages (chelifores) that allow them to manipulate food items before consumption, some species lack these limbs altogether and instead rely on the moveable elongate proboscis for suction feeding (Fig. 3E, H). Chelifore-less pycnogonids, such as *Pycnogonum,* ammotheids and aschorhynchids, are known to consume soft-bodied and gelatinous organisms including actinians, hydroids and even pelagic ctenophores [33, 34] (Fig. 3H). The lumen of the pycnogonid proboscis is voluminous and directly connects proximally with a pharyngeal filtering system (oyster basket) which consists of dense packed cuticular denticles and bristles that are used to process ingested solid particles [35] (Fig. 3G). Pycnogonids also have anterior palps and/or setae with a sensorial function that allow them to navigate the environment and locate food items. Thus, *H. sparsa* and pycnogonids represent approximate ecological analogues that share a similar lifestyle (slow moving marine epibenthic), overall body construction (tubular cuticular trunk with mostly homonomous elongate clawed walking legs) and sensorial appendages (anterior palps or tentacles). These taxa also share similar oral adaptations for feeding (Fig. 3). The elongate bulbous head of *H. sparsa* resembles the moveable pycnogonid proboscis (Fig. 3A, B, E), and in both cases the terminal mouth is followed by an enlarged cavity expressed as the buccal chamber [18] and the pharynx respectively [35] (Fig. 3B, G). The presence of sclerotized acicular denticles on the foregut of *H. sparsa* [18] (Fig. 3C, D) evokes the cuticular structures that conform the pharyngeal filter system of pycnogonids in terms of their organization (small and tightly packed sclerotized elements) and location [33, 35] (Fig. 3G). The fact that pycnogonid clutters have been recorded consuming pelagic ctenophores and other gelatinous organisms [34] (Fig. 3H) adds further behavioural feeding parallels with *H. sparsa* as informed by MCZ.IP.102104 (Fig. 1, 2). It is worth noting that carnivorous pycnogonids are technically classified as parasites since they can feed off living organisms without killing them [33]. The available fossil data is insufficient to confidently determine whether *H. sparsa* was an obligate scavenger that exclusively fed on carcasses, or a facultative parasite and opportunistic scavenger akin to extant pycnogonids.

**Figure 3.**
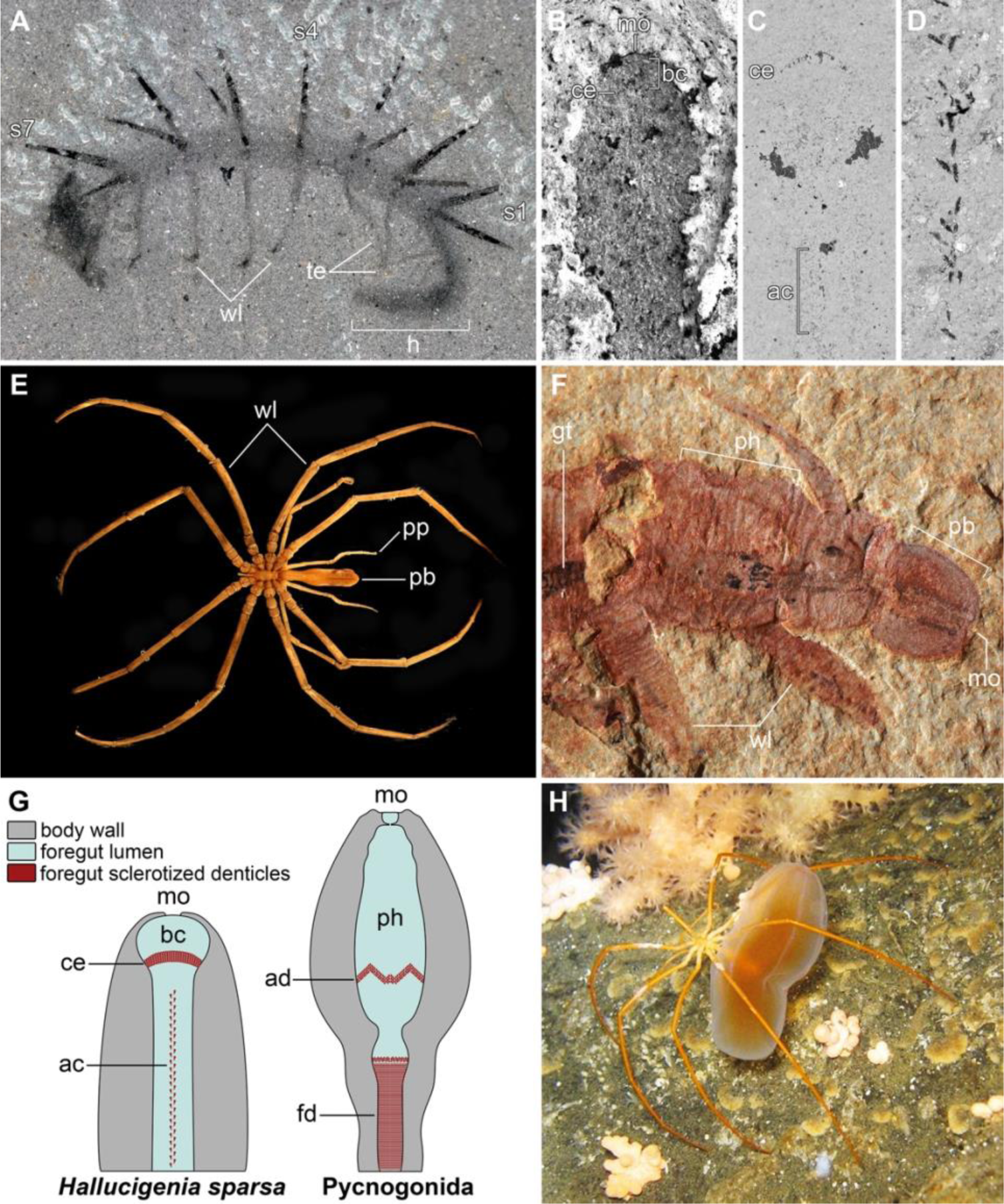
Morphological and ecological adaptations for suction feeding in Cambrian armoured lobopodians and extant pycnogonids. **A.** NMNH-198658, overview of *Hallucigenia sparsa* in lateral view showing elongate bulbous head region and sensorial anterior tentacles [18]. **B.** ROM 63146, scanning electron micrograph of head region in *H. sparsa* in dorsal view showing details of oral region [18]. **C.** ROM 63146, backscattered electron micrograph of magnified head region in *H. sparsa* showing buccal chamber and carbonaceous preservation of circumoral sclerotized elements and sclerotized aciculate denticles in foregut [18]. **D.** Magnification of sclerotized aciculate denticle row in foregut [18]. **E.** Overview of extant chelifore-less pycnogonid *Colossendeis proboscidea* showing elongate proboscis and sensorial pedipalps (image copyright held by The Cole Museum of Zoology, University of Reading, UK). **F.** ELEL-SJ101888A, anterior morphology of the lobopodian *Onychodictyon ferox* showing robust proboscis with terminal mouth and well-developed pharynx connected to straight gut tract [41]. **G.** Schematic comparison between the proposed functional analogues in the head of *Hallucigenia sparsa* and the proboscis of extant pycnogonids (modified from ref. [18, 35]). **H.** Individual of *Colossendeis megalonyx* consuming a pelagic beroid ctenophore (reproduced from ref. [34]. Abbreviations: ac, acicular denticles; ad, anterior denticles; bc, buccal cavity; ce, circumoral elements; fd, filtering denticles; h, head; mo, mouth opening; ph, pharynx; pb, proboscis; pp, pedipalps; s*n,* dorsal spines; te, tentacles; wl, walking legs.

A second key insight is that all *H. sparsa* on MCZ.IP.102104 appear to be contemporaneous in terms of their ontogeny based on their similar size, and the fact that the seven complete individuals are evenly spaced throughout the *Xanioascus* carcass. The taphonomic evidence supporting *in situ* burial for the fossil assemblage (see above) argue that the spatial distribution of *H. sparsa* is not the result of size sorting due to transport. The presence of several small (possibly juvenile) individuals of *H. sparsa* feeding on the *Xanioascus* carcass simultaneously is consistent with a swarm-like behaviour during foraging. Indeed, the lobopodians are spread out non-randomly as if to maximise feeding area while minimizing direct intraspecific competition. Swarming behaviour is well documented in modern marine ecosystems and is particularly relevant in the case of opportunistic scavenger communities of epibenthic invertebrates that rely on episodic carcass falls such as whales [36] and jellyfish blooms [37]. Whilst expressed at the much smaller scale of the *Xanioascus* carcass, MCZ.IP.102104 shows that *H. sparsa* engaged in collective scavenging feeding behaviour. A strikingly similar example is known from the early Cambrian Chengjiang biota of South China, in which several small specimens of the armored lobopodian *Microdictyon sinicum* have been observed in direct association with the soft-bodied organism *Eldonia* [38]. Initially interpreted as an unconventional pseudopelagic lifestyle for *M. sinicum* based on the view of *Eldonia* as a pelagic organism [38], *Eldonia* is now regarded as a soft-bodied epibenthic stalked deuterostome [39]. These associations have been subsequently reinterpreted as evidence of *M. sinicum* feeding on *Eldonia* carcasses on the seafloor [17]. Another relevant case from Chengjiang consists of dense mass associations of palaeoscolecids and lobopodians interpreted as opportunistic feeding swarms around nutrient rich patches such as animal carcasses [17]. However, these differ from MCZ.IP.102104 in that there is no clear evidence of a discrete carcass serving as the food source. A last comparable demonstration of swarm scavenging comes from the Burgess Shale, as an assemblage of at least five individuals of the archaeopriapulid worm *Ottoia prolifica* (arguably) devouring the carcass of the large euarthropod *Sidneyia inexpectans* [1, 10]. Opportunistic feeding swarms are commonplace in benthic ecosystems and support diverse animal communities in otherwise macronutrient depleted local environments. Epibenthic scavengers are terrifically effective bioturbators, with empirical data showing that an entire jellyfish carcass may be consumed within 2.5 hours in modern marine ecosystems [37]. Thus, direct evidence of this ephemerous ecological phenomenon is extremely rare in the fossil record. The new data on MCZ.IP.102104 provides an rare glimpse into the intricate trophic ecology of the middle Cambrian seafloor.

### The ecological role of Cambrian lobopodians

The results of this study cast new light on the enigmatic ecology and feeding behaviour of *Hallucigenia sparsa* from the mid-Cambrian Burgess Shale as an opportunistic and gregarious suction feeding scavenger of gelatinous (and likely more generally soft-bodied) food items. It is notable that most well-known lobopodians within the stem lineage of Onychophora (e.g. *Cardiodyction, Microdictyon, Onychodictyon,* other *Hallucigenia* [05, 18, 25, 38, 40]) closely resemble *H. sparsa* in having a tubular body with metameric sclerotized dorsal armature, modified tentacle-like anterior appendages, elongate bulbous heads with a terminal mouth but without external mouthparts, and simple digestive tracts [08, 16, 20] (Fig. S3). Indeed, some lobopodians have even less differentiated appendages (e.g. *Paucipodia* [41), or feature different types of dorsal epidermal armour (e.g. *Diania* [42]), yet consistently feature bulbous heads with terminal mouths without obvious external mouthparts and simple gut tracts. These morphological parallels are particularly relevant when considering that *M. sinicum* and *H. sparsa* also share behavioural evidence for swarming on soft-bodied carcasses that point to a fundamentally similar feeding ecology [17, 38] (Fig. 1, 2; Fig. S3). *Onychodictyon ferox* is exceptionally informative in this regard thanks to its robust proboscis with a terminal mouth opening coupled with a well-developed muscular pharynx and regarded as having a suction feeding function [43] (Fig. 3F; Fig. S3D). Even the highly derived luolishaniids, which have elaborate setulose anterior trunk appendages and variable degrees of dorsal armour ranging from exuberant to non-existent [05, 29, 44, 45, 46], must have employed some degree of suction feeding to ingest the suspended organic matter sieved by their limbs from the water column. The fact that aciculate sclerotized denticles have been reported in the mouth of the luolishaniid *Ovatiovermis cribatus* [29], closely resembling those of *H. sparsa* [18], adds another line of evidence pointing to adaptations for suction feeding. These ecological interpretations of the functional morphology of small bodied lobopodians, based on the present data on *H. sparsa*, suggest that suction feeding was ancestral for total-group Onychophora (Fig. 4). The origin of an active macropredatory mode of life that typifies modern onychophorans [47], including the presence of internalized paired jaws from modified walking legs and slime-shooting papillae to immobilize prey, most likely evolved after the terrestrialization of this clade during the mid to late Paleozoic [48, 49]. This phylogenetic pattern contrasts with the feeding ecology of lobopodians within the euarthropod stem lineage (lower stem-group Euarthropoda [50]), which showcase clear adaptations for macrophagy and predatory lifestyles including anterior raptorial appendages, circumoral plates, pharyngeal teeth, and complex mid-gut glands [17, 18, 51, 52, 53, 54]. The evolution of paired body flaps in these large-bodied lobopodians also greatly enhanced their mobility and swimming capabilities, consistent with an active macropredatory behaviour [17, 49, 50, 51, 52, 55]. Thus, the main Cambrian diversification of Panarthropoda was accompanied by a fundamental ecological partition (Fig. 4). The euarthropod stem lineage was dominated by large bodied lobopodians that occupied a primary role as active macrophagous predators and scavengers, including both epibenthic and nektobenthic forms. By contrast, the onychophoran stem lineage evolved as strictly epibenthic small bodied (typically armoured) lobopodians specialized in suctorial strategies ranging from fluid scavenging soft-bodied carcasses (*Hallucigenia, Microdictyon*) to suspension feeding in the water column (luolishaniids). *H. sparsa* and other small-bodied (typically armoured) lobopodians would have played a substantial scavenging trophic role in Cambrian ecosystems by breaking down carcasses of gelatinous and other soft-bodied organisms, thereby facilitating the reintegration of organic carbon and energy into the environment. The new evidence on *H. sparsa* further demonstrates that lobopodians directly contributed to the transfer of energy from the plankton to the benthos due to their opportunistic consumption of pelagic organisms such as comb jellies (Fig. 1, 2), as also observed in extant pycnogonids [38] (Fig. 3H). Consequently, this ecological strategy helped prevent the excessive accumulation of gelatinous and soft bodied carcass material that could lead to ecosystem fouling, which has been shown to the essential for maintaining viable epibenthic communities in modern oceans [36, 37]. Densely fossil assemblages of *Hallucigenia* [18] and other small-bodied lobopodians [17] offer a glimpse into the abundance of these animals in the Cambrian seafloor and their ecological contribution to marine ecosystems millions of years before the evolution of modern swarm scavengers such as decapod crustaceans.

**Figure 4.**
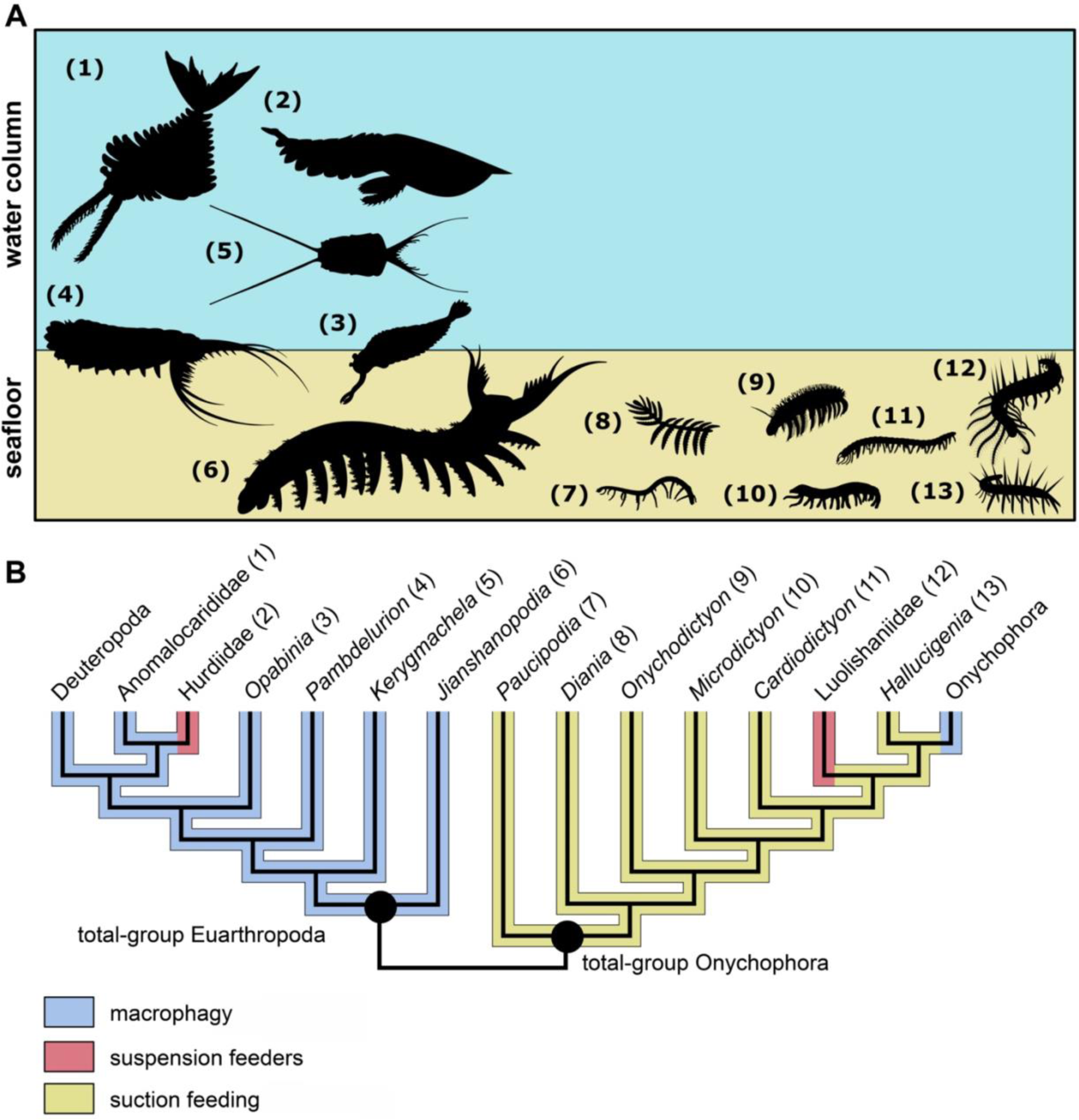
Major phylogenetic and ecological partition of Cambrian lobopodians and panarthropod phyla. **A.** Lobopodians with raptorial appendages and external mouthparts are typically large bodied (>10 cm body length) and have a nektobenthic lifestyle, whereas armoured lobopodians with elongate bulbous heads are typically small bodied (<10 cm body length) and are exclusively epibenthic; silhouettes sourced from PhyloPic. **B.** Simplified phylogenetic relationships of Panarthropoda showing proposed distribution of feeding strategies in total-group Euarthropoda and total-group Onychophora; topology based on refs [5, 18, 25].

**Figure 5.**
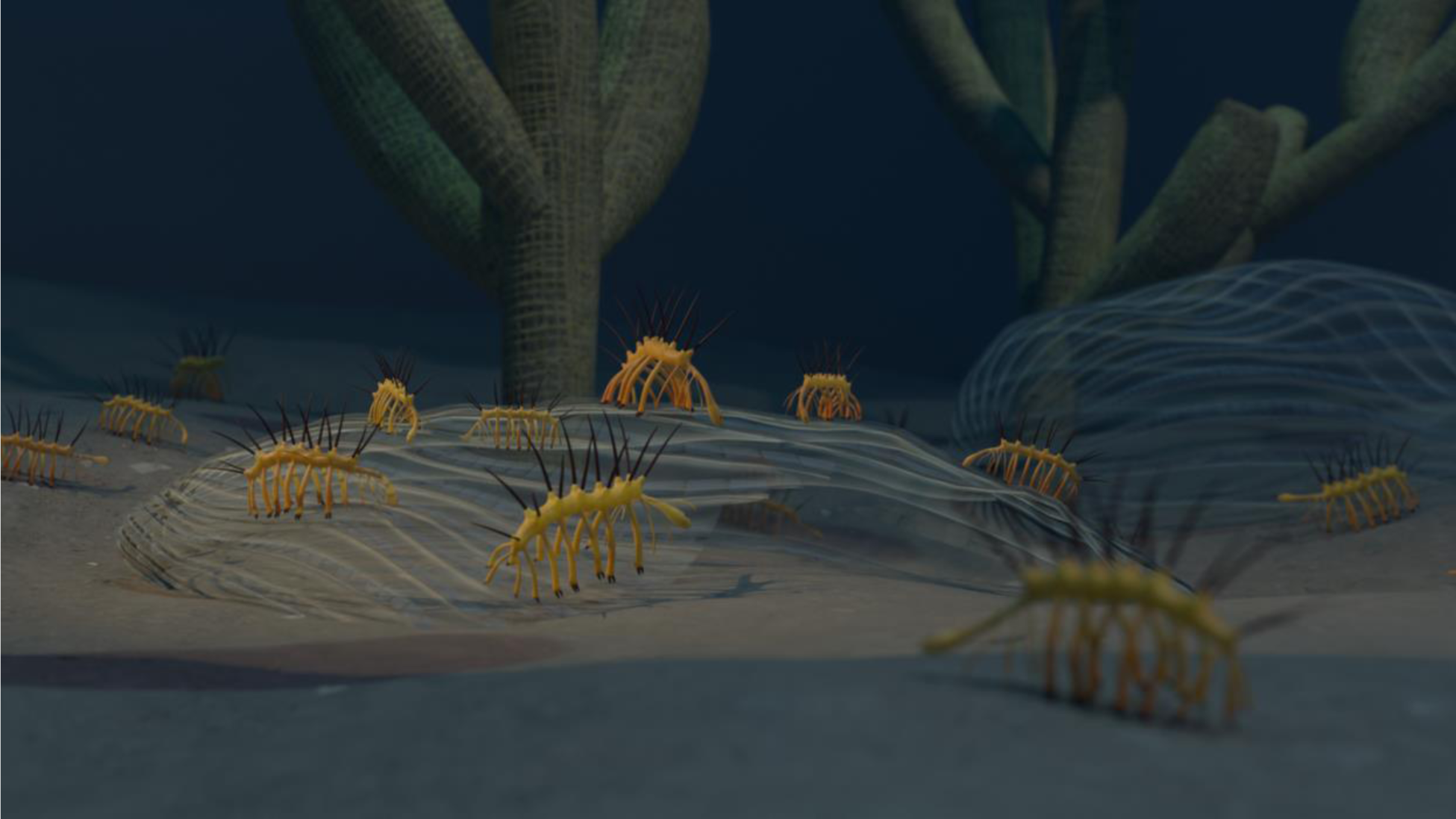
Artistic reconstruction of *Hallucigenia* swarm feeding on *Xanioascus* carcasses. Created by Franz Anthony.

## Methods

The studied material is housed at the Invertebrate Paleontology collections at the Museum of Comparative Zoology at Harvard University (MCZ.IP). Fossils were photographed under cross polarized illumination, both dry and wet (70% ethanol), with a Nikon Z8 (Full Frame, 48 MP) fitted with a Macro Lens Nikkor Z MC 50 mm. Digital photographs were stacked with the software Zerene Stacker. Figures were drafted with Adobe Photoshop 2023 and Inkscape.

## Data availability

MCZ.IP.102104 is housed at the Invertebrate Paleontology Collection of the Museum of Comparative Zoology, Harvard University, USA. All data associated with this study is available in the manuscript and supplementary information.

## Acknowledgments

Thanks to Jessica Cundiff (Museum of Comparative Zoology, Harvard University) for facilitating access to the studied material. Thanks to Bruno Becker Kerber for assistance with photographic hardware and software. Martin Smith (Durham University) generously shared digital images of *Hallucigenia sparsa*; figured material deposited at the National Museum of Natural History, Washington D.C., USA (NMNH), and the Royal Ontario Museum, Toronto, Canada (ROM). Ailin Chen (Yuxi Normal University) generously shared digital images of *Microdictyon sinicum*; figured material deposited at the Yuxi Normal University Research Center of Paleobiology (YRCP). Qiang Ou (China University of Geosciences) generously shared digital images of *Onychodictyon ferox*; figured material deposited at the Early Life Evolution Laboratory, China University of Geosciences, Beijing, China (ELEL). Jean-Bernard Caron (Royal Ontario Museum) generously shared digital images of *Xanioascus canadensis* and *Ctenorhabdotus capulus*; figured material deposited at the Royal Ontario Museum (ROM). The Cole Museum of Zoology (University of Reading) generously granted permission to use the digital photograph of *Colossendeis proboscidea*.

## Ethic declarations

The author declares no competing interests.

---

**Figure S1.**
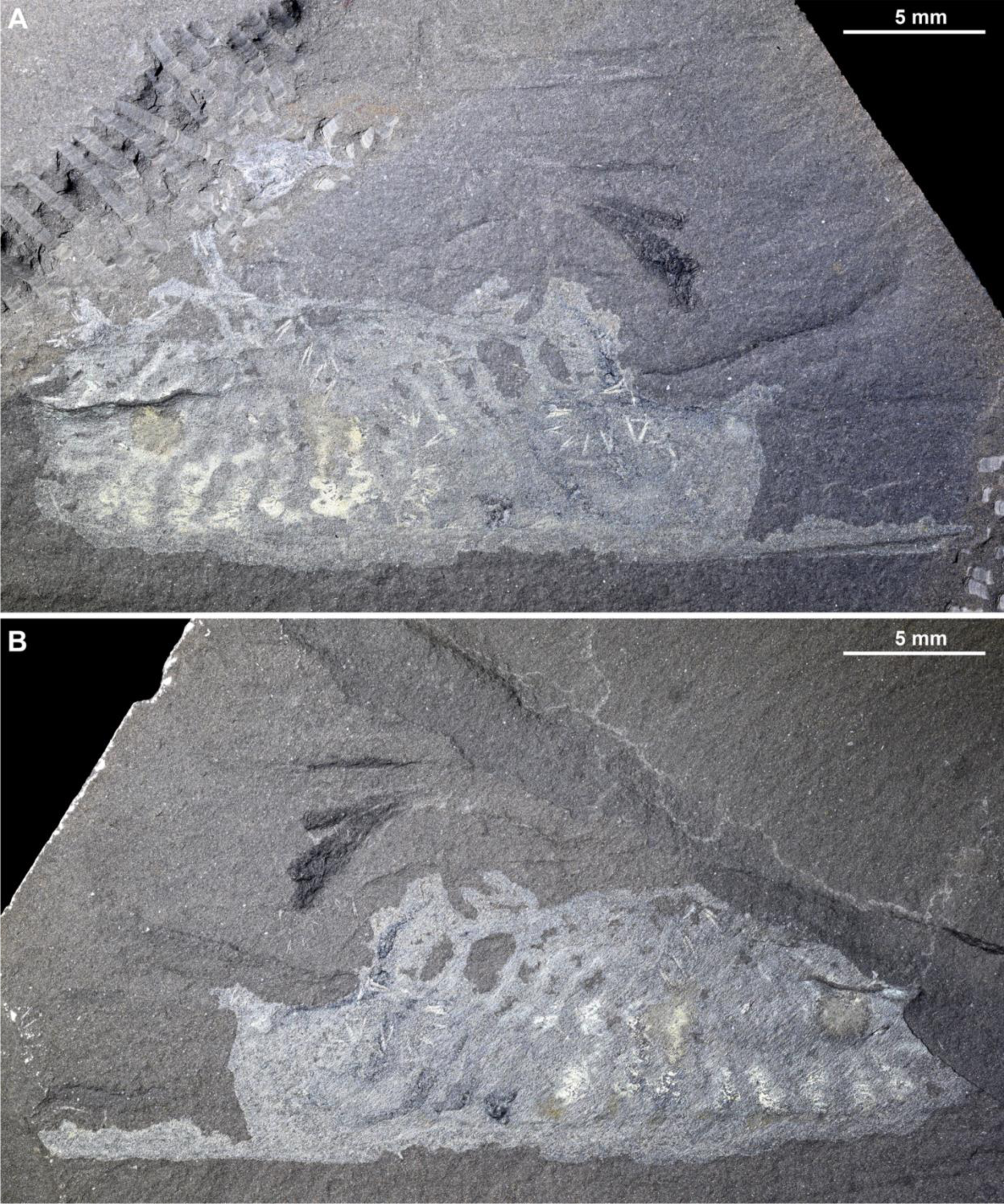
Unannotated digital images of MCZ.IP.102104 photographed wet under cross polarized illumination emphasizing highly reflective carbonaceous films. **A**. Part; see also Fig. 1A. **B**. Counterpart; see also Fig. 2A.

**Figure S2.**
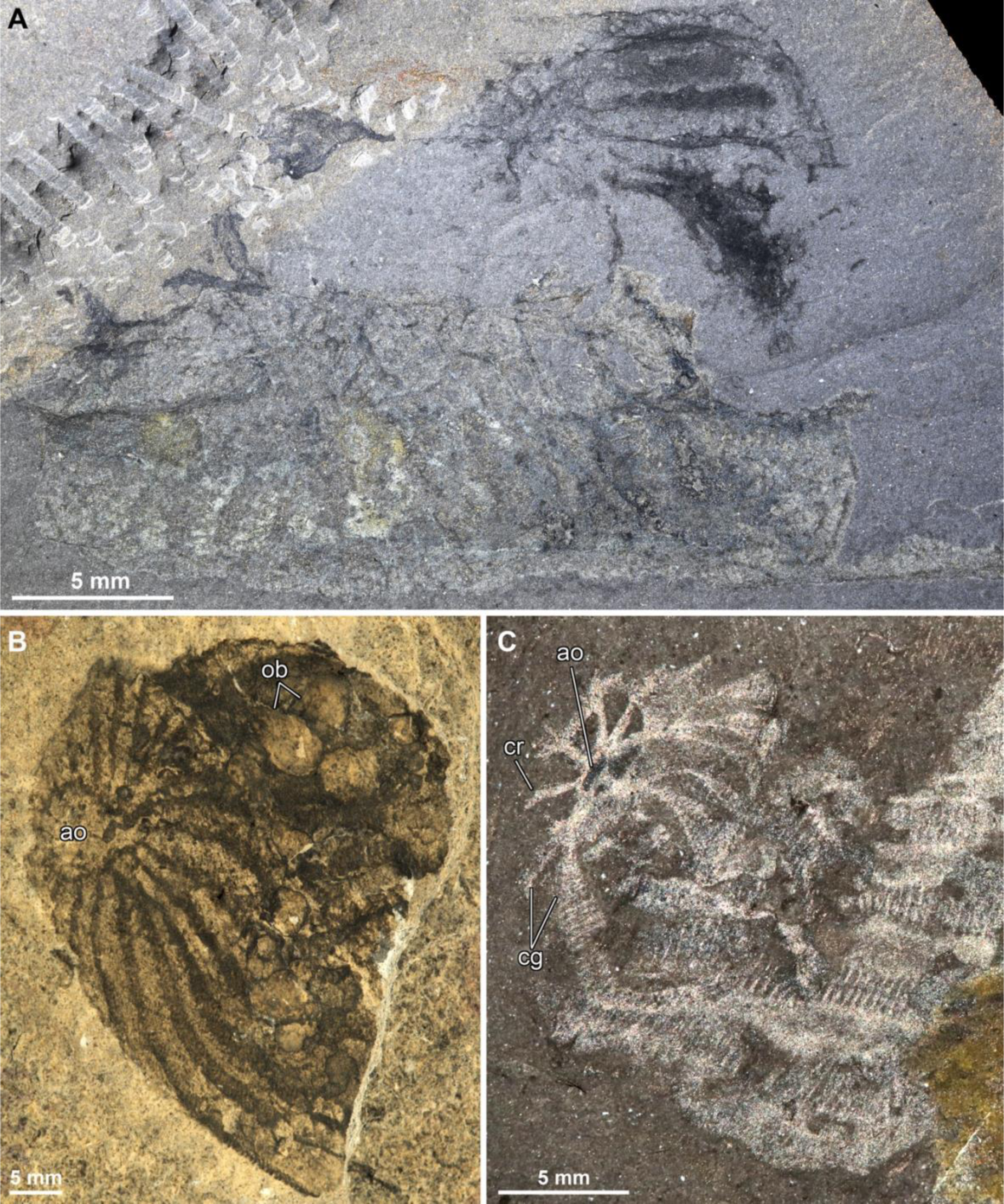
Morphological comparison between pelagic ctenophores from the middle Cambrian Burgess Shale. **A.** MCZ.IP.102104 part, unannotated digital photograph of soft-bodied ctenophore carcass tentatively interpreted as *Xanioascus* showing evidence of comb rows, ciliary grooves, an apical organ without a capsule and internal ovoid bodies; see also Fig. 1B. **B.** Well preserved individual of *Xanioascus canadensis* showing apical region, comb rows and internal ovoid bodies (ROM 43191; image courtesy of Jean-Bernard Caron). **C.** Well preserved aboral region of *Ctenorhabdotus capulus* showing apical organ connected with radial ciliary grooves and comb rows (ROM 51439; image courtesy of Jean-Bernard Caron). Abbreviations: ao, apical organ; cg, ciliary grooves; ct, comb rows; ob, ovoid bodies.

**Figure S3.**
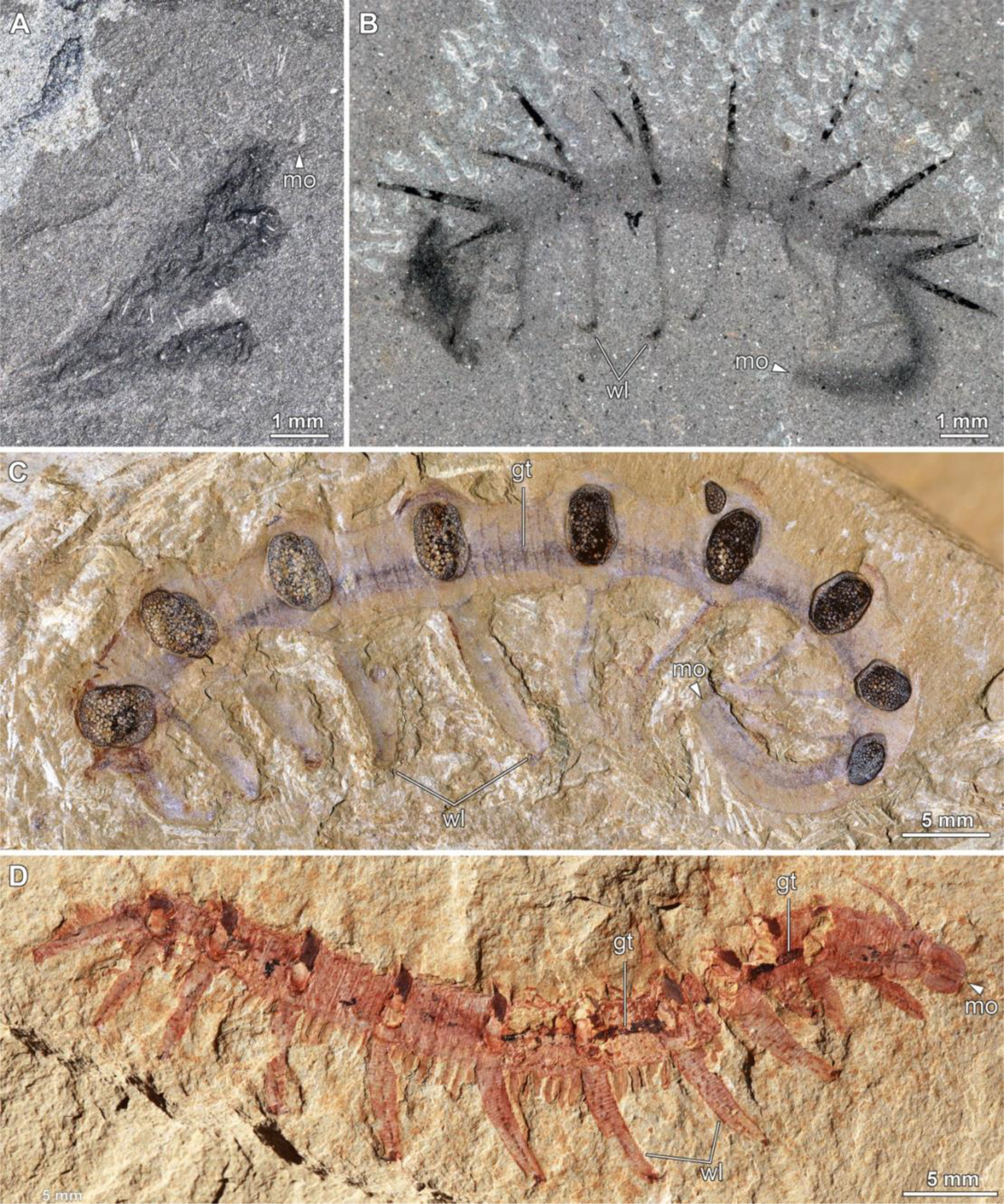
Morphological organization of Cambrian small-bodied armoured lobopodians. **A.** Small complete individual of *Hallucigenia sparsa* (MCZ.IP.102104 counterpart) in lateral view showing elongate bulbous head region; see also Figure 2C. **B.** Complete individual of *Hallucigenia sparsa* (NMNH-198658; image courtesy of Martin R. Smith) in lateral view showing elongate bulbous head region with terminal mouth opening; see ref. [18]. **C.** Complete individual of *Microdictyon sinicum* (YRCP0041; image courtesy of Ailin Chen) in lateral view showing elongate bulbous head region with terminal mouth opening and simple gut tract throughout the body; see ref. [40]. **D.** Complete individual of *Onychodictyon ferox* (ELEL-SJ101888A); image courtesy of Qiang Ou) in lateral view showing robust proboscis and terminal mouth, well developed pharynx and simple gut tract throughout the body; see ref. [43]. Abbreviations: gt, gut tract; mo, mouth opening; wl, walking legs.

